# The Centrosome Controls the Number and Spatial Distribution of Microtubules by Negatively Regulating Alternative MTOCs

**DOI:** 10.1101/153304

**Authors:** Maria P. Gavilan, Pablo Gandolfo, Fernando R. Balestra, Francisco Arias, Michel Bornens, Rosa M. Rios

## Abstract

In this work, we have investigated in mammalian cells how microtubule nucleation at centrosome and Golgi apparatus are coordinated, using genetic ablation of three γ-TuRC binding proteins -AKAP450, Pericentrin and CDK5Rap2- and the PLK4 inhibitor centrinone. We show that centrosomal microtubule nucleation is independent of Golgi activity whereas the converse is not true: nucleation on the Golgi negatively correlates with the number of centrosomes. In addition, depleting AKAP450 in cells lacking centrioles, that abolishes Golgi nucleation activity, leads to microtubule nucleation from numerous cytoplasmic Golgi-unbound acentriolar structures containing Pericentrin, CDK5Rap2 and y-tubulin. Strikingly, centrosome-less cells display twice higher microtubule density than normal cells, suggesting that the centrosome controls the spatial distribution of microtubules, not only by nucleating them, but also by acting as a negative regulator of alternative MTOCs. Collectively, the data reveals a hierarchical control of microtubule nucleation, with the centrosome regulating this process in a more complex manner than usually thought. It also unveils mechanisms that could help understanding MT network reorganization during cell differentiation.

---

The centrosome is the major microtubule (MT)-organizing center (MTOC) of animal cells. It is formed of a pair of centrioles organizing a matrix, the pericentriolar material (PCM), were MT nucleation and anchoring activities are localized [1]. In the last years, it has been convincingly shown that MT nucleation also takes place in other subcellular locations [2, 3]. During interphase, the most important additional MTOC is the Golgi Apparatus (GA) that has been estimated to contribute almost 50% of the total MTs in RPE1 cells [4, 5]. Throughout mitosis, spindle MTs and kinetochores are also active non-centrosomal MTOCs [6, 7].

MT nucleation primarily relies on γ-tubulin and its associated proteins (GCP2-GCP6). In higher eukaryotes, these proteins form ring-shaped complexes known as γ-TuRCs that serve as scaffolds for tubulin dimers in order to promote MT polymerization [8]. In addition, efficient γ-tubulin mediated MT nucleation depends on additional regulatory factors such as Mozart1/MZT1 or NEDD1 that activate or target γ-TuRCs to MTOCs [2, 9].

The centrosomal proteins Pericentrin (PCNT), AKAP450 (also known as AKAP350 or CG-NAP), CDK5Rap2 (Cep215) and myomegalin contain conserved, yet degenerate, motifs for γ-TuRCs binding [10] and they have been shown to bind γ-TuRC in vivo [11-14]. Furthermore, PCNT interacts with AKAP450, and both proteins contribute to CDK5Rap2 and myomegalin recruitment to the centrosome [15-18]. Thus, proteins acting as γ-TuRC receptors seem to assemble into higher-order complexes, which might ensure efficient γ-TuRC recruitment to the centrosome. Based on these features, a role of these proteins in MT nucleation at the centrosome has been generally assumed. However, direct evidence supporting this view is scarce and mechanisms regulating centrosomal MT nucleation, especially during interphase, remain poorly understood.

With the exception of PCNT, which has only been detected at the GA of skeletal muscle fibers, the three other γ-TuRC-binding proteins mentioned above, i.e., AKAP450, CDK5Rap2 and myomegalin, do also localize at the cis-face of the GA [16-19]. The *cis*-Golgi protein GM130 recruits AKAP450, which in turn, recruits both CDK5Rap2 and myomegalin [16, 17]. AKAP450 has been proven to be the major regulator of MT nucleation at the GA as either si-RNA driven knocking-down, gene knock-out or dissociation of AKAP450 from the GA completely abolishes Golgi-associated MT nucleation [19-21]. CDK5Rap2 and myomegalin appear to only facilitate the process by providing MT stabilization/anchoring activities [14, 21]. In addition to these players, assembly of Golgi-associated MTs requires the dynein/dynactin complex [19, 20] and the MT-binding proteins CLASPs [4]. Other proteins such as Microtubule-Crosslinking Factor 1 (MTCL1) that binds both AKAP450 and CLASPs [22], and Calmodulin-Regulated Spectrin-Associated Protein Family Member 2 (CAMSAP2) that binds myomegalin 8 [21] also appear to play a role in the dynamics of Golgi-nucleated MTs.

How MT nucleation associated to two different organelles is controlled and coordinated is an open question. Indeed, it is known that the respective contribution of both organelles to the overall MT network organization vary along the cell cycle or during cell differentiation. For instance, during G2-M transition, centrosome maturation correlates with a gain of its MT nucleation activity [9], while MT-nucleation activity of the GA is lost [23]. The process is reversed after mitotic exit when GA MT nucleating activity is fully recovered. Increase of PCM size and enhanced MT nucleation have been also reported in response to pro-inflammatory stimuli [24]. On the contrary, MT nucleating activity of the centrosome is down-regulated, or even completely abolished, during cell differentiation [25-28]. These functional changes are usually linked to variations in PCM levels and/or shedding of some PCM components. However, the underlying molecular mechanisms regulating these processes have still to be discovered. Even less is known about how the GA regulates its MT nucleating activity. Elucidating the regulation of MT nucleation activities of the centrosome and the GA is critical to understand how functions of both organelles are orchestrated in order to create an appropriate MT array. For instance, MT nucleation activity of the GA seems to increase in cells lacking centrosomes [21]. To gain further insights into this regulation, we have manipulated MT nucleation activities of both the centrosome and the GA in hTERT-RPE1 cell lines depleted of AKAP450, PCNT and CDK5Rap2 using centrinone which generates cells lacking centrioles [29].

We report here the existence of two kinds of γ-TuRC-binding protein complexes: those containing AKAP450 that localize at both the GA and the centrosome, and those containing PCNT that specifically accumulate at the centrosome. These complexes exhibit antagonistic effects on GA-associated MT nucleation while they do not apparently participate in centrosomal MT nucleation. In the absence of centrosome, PCNT-based complexes redistribute from the PCM to the GA. When the centrosome is eliminated and GA-associated MT nucleation abolished by AKAP450 depletion, PCNT-based complexes become competent for MT nucleation in the cytoplasm where they behave as acentriolar MTOCs. The consequences of these manipulations on global MT network organization have been quantified. Collectively, our data leads to an unexpected vision of how the centrosome can control the spatial organization of the MT network in cells.

## RESULTS

### Characterization of akap9, cdk5rap2 and pcnt knock-out hTERT-RPE1 cell lines

To generate AKAP450, CDK5Rap2 and PCNT B/kendrin (PCNT hereafter) hTERT-RPE1 knock-out (KO) cell lines we used CRISPr/Cas9 nickase-mediated mutagenesis. In order to improve efficiency and specificity of the process, we developed a vector containing two cloning sites for two single guide RNAs (sgRNAs; see Fig S1A and Material and Methods for details). The sgRNAs were designed to target the first exons of each protein (exon 2, 1 and 5 for *akap9*, *cdk5rap2* and *PCNT* genes respectively, Fig. S1B), in order to prevent the presence of truncated proteins in mutant cell lines. KO clones for each gene were initially identified by immunofluorescence (IF) and, after expansion, characterized by western-blotting (WB) and IF using several antibodies as indicated (see schemes in Fig. 1A, D and F). All selected clones bear alleles with premature stop codons as a consequence of the introduced mutations (Fig. S2). The amino-acid sequence of putative truncated proteins expressed in KO clones, if any, are depicted in Fig. S2A-C. Notably, cell viability and proliferation were not compromised in any of the KO cell lines.

**Figure 1.**
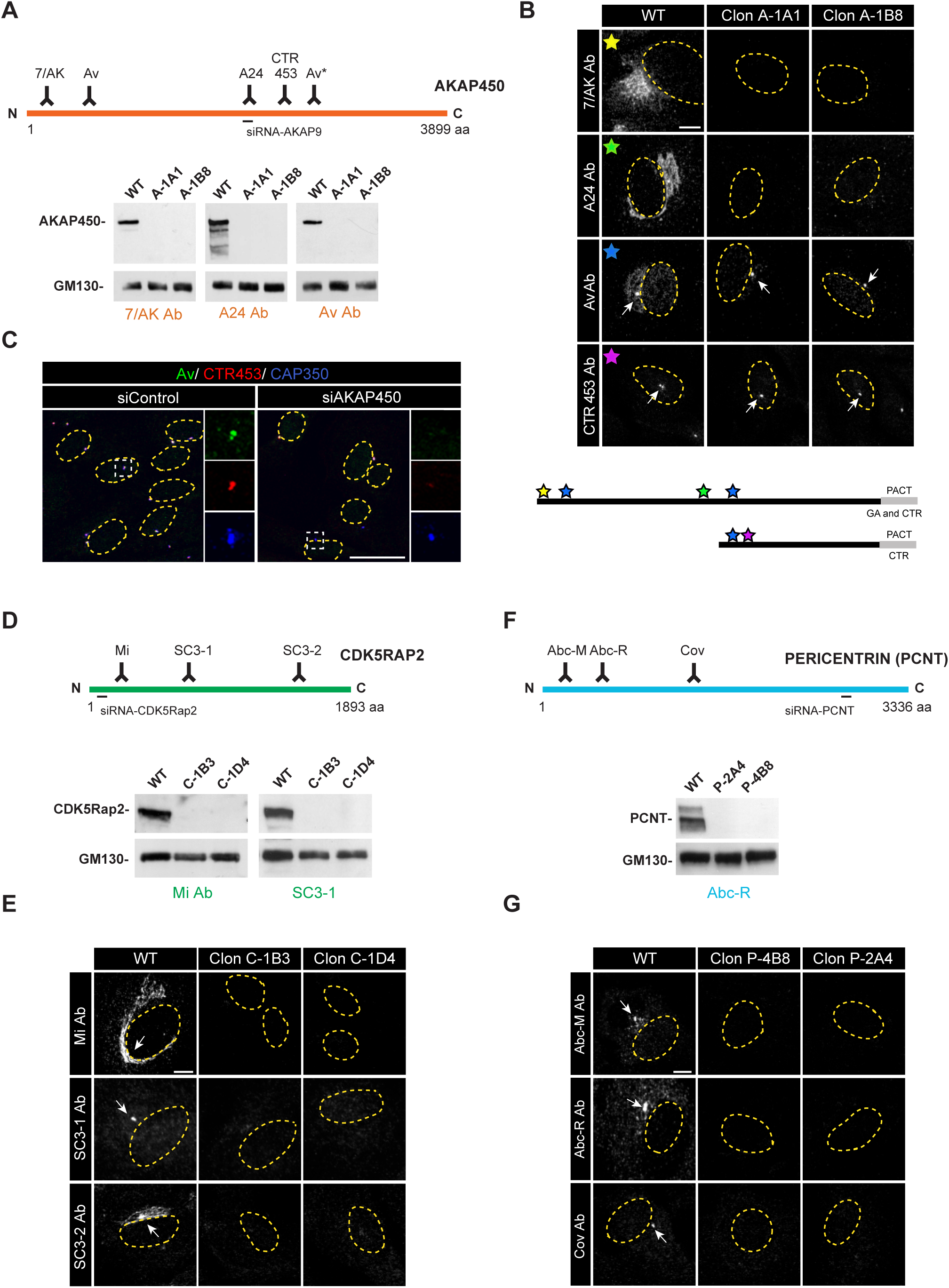
Characterization of *akap9*, *cdk5rap2* and *pcnt* knock-out cell lines. (**A**) (Top) Schematic representation of AKAP450 protein illustrating localization of the epitopes recognized by the antibodies used in this study and the sequence targeted by the siRNA used in (C). (Bottom) Representative Western blot (WB) of RPE-1 control cells and the two selected *akap9* mutated clones (A-1A1 and A-1B8) probed with three anti-AKAP450 antibodies (A24, 7/AK and Av) as indicated. GM130 was used as a loading control. (**B**) (Top) Confocal IF images of RPE-1 control cells and RPE-1 *akap9* mutated clones with different anti-AKAP450 antibodies as indicated. Each antibody is represented by a coloured star. Centrosomes are indicated by white arrows. The yellow dashed line indicates the nucleus contour. Note that a centrosomal signal persists when Av and CTR453 antibodies are used. (Bottom) Putative AKAP450 isoforms and localization of the epitopes recognized by each antibody. (**C**) RPE-1 cells transfected with either control (left) or AKAP450 (right) siRNAs and triple labeled with Av (green), CTR453 (red) and anti-CAP350 (blue, as a centrosomal marker) antibodies. Merged images are shown. High magnification images of centrosomes in boxed areas are shown at right with single labelings. (**D**) (Top) Same as in (A) but for CDK5Rap2 protein. (Bottom) WB of RPE-1 control cells and *cdk5rap2* mutated clones (C-1B3 and C-1D4) tested with two anti-CDK5Rap2 antibodies as indicated. (**E**) IF images of RPE-1 control cells and RPE-1 *cdk5rap2* mutated clones stained with the three available anti-CDK5Rap2 antibodies. (**F**) (Top) Schematic diagram of the PCNT protein including information of both siRNA-targeted sequence and antibody-recognized regions as in (A) and (D). (Bottom) WB analysis of RPE-1 *pcnt* mutated clones. (**G**) RPE-1 control cells and *pcnt* mutated clones (P-2A4 and P-4B8) were analyzed by IF with different anti-PCNT antibodies as depicted. Scale bars= 5 μm (B, E, G), 25 μm (C).

Four anti-AKAP450 antibodies recognizing different epitopes all over the protein were used (Fig. 1A). Three of them, i.e. 7/AK, A24 and Av antibodies, labeled both the GA and the centrosome in wild-type (WT) cells (Fig. 1B), although the GA labeling with Av antibody was very weak (Fig. 1B). On the other hand, the monoclonal antibody CTR453 only decorated the centrosome. By WB, none of the antibodies tested (CTR453 antibody does not work for WB) revealed a signal in extracts of the *akap9* KO clones suggesting that selected clones did not express any *akap*9 gene product (Fig. 1A). By IF, GA and centrosome labeling displayed by A24 and 7/AK antibodies as well as the weak GA staining displayed by the Av antibody disappeared in *akap9* KO cells (Fig. 1B). However, centrosomal labeling persisted with both the Av and the CTR453 antibodies. These results suggested that, although the major AKAP450 isoform is absent in *akap9* KO cells, minor centrosome-specific isoform/s is/are still present in *akap9* KO cells. Our data are compatible with this centrosomal isoform consisting of the C-terminal middle part of the protein. Indeed, such an isoform would not be recognized by antibodies against the N-terminal end of the protein and should not be targeted to the GA. However, Av Ab results did not entirely fit with this possibility since it was raised against the N-terminal part of the protein. To clarify this issue, we further characterized Av antibody in transiently transfected RPE-1 cells expressing different fragments of AKAP450 fused to GFP (Fig. S3A). Av Ab recognized both AK1 (1-1004 aas) and AK3 (1708-2864) fragments (Fig. S3B). Interestingly, the AK3 fragment contains the exon 29 that is recognized by CTR453 [30]. Finally, we performed siRNA experiments targeting *akap9* mRNA (Fig. 1C). siRNA reduced the centrosomal pool of AKAP450 recognized by both Av and CTR453 antibodies in a significant percentage of cells, which demonstrated that the centrosomal staining was AKAP450 specific (Fig. 1C). In summary, our data shows that *akap*9 KO cell lines lack AKAP450 but maintain a minor centrosomal pool, undetectable in WB of whole cell extracts, which is only revealed by using specific antibodies. A predicted AKAP450 isoform containing exons 27 to 50 has been reported that fits with our results (Uniprot ref. number H7BYL6). Hereafter, we refer to this minor centrosomal AKAP450 as AKAP450^cent^, to distinguish it from the major AKAP450 population, referred to as AKAP450.

To analyze *cdk5rap2* KO clones we employed three antibodies recognizing different domains of the protein (Mi, SC3-1, SC3-2; Fig. 1D). Polyclonal SC3-1 and SC3-2 antibodies were generated during this study (see M&M). The three antibodies decorated both the GA and the centrosome, although with a wide range of intensities for each organelle. Thus, while Mi Ab strongly recognized the GA, the SC3-1 antibody almost exclusively labeled the centrosome. None of the three antibodies detected CDK5Rap2 by IF in KO clones (Fig. 1E). In addition, Mi and SC3-1 antibodies did not reveal any signal when tested by WB on selected KO cell extracts suggesting that we were most probably dealing with a full CDK5Rap2 KO cell line. Finally, we carried out similar experiments to characterize *pcnt* KO cell lines by using three different antibodies as represented in Fig. 1F. The three antibodies stained essentially the centrosome, with a weak spotty pericentrosomal staining with Abc-M and –R Abs. No IF signal was observed with any of the antibodies tested in the selected KO clones (Fig. 1G) nor with the only anti-PCNT antibody that worked in WB analysis. Given the N-terminal distribution of epitopes recognized by the available anti-PCNT antibodies, the possibility that some PCNT C-terminal isoform/s are still expressed in KO clones, like for AKAP450^cent^, cannot be excluded. However, we did not find any evidence for such a possibility in database.

Since AKAP450, PCNT and CDK5Rap2 have been shown to interact *in vivo*, we wondered whether the absence of one of them could affect the protein level of the other ones. To answer this question, we analyzed the expression levels of the three proteins in all KO cell lines. As shown in Fig. S3C, the expression level of each of these proteins was not affected by depletion of the other two proteins.

### Distribution of AKAP450, AKAP450^cent^, CDK5Rap2 and PCNT at the GA and the centrosome

In order to clarify the relationship between AKAP450, AKAP450^cent^, CDK5Rap2 and PCNT proteins, we analyzed and quantified by IF their respective distributions at both the centrosome and the GA in either WT, *akap9, cdk5rap2* and *pcnt* KO cells. As the GA usually surrounds the centrosome, we performed these experiments in nocodazole (NZ)-treated cells since MT disassembly induces fragmentation and dispersion of the Golgi ribbon and allows better discrimination of centrosome- and GA-associated protein pools.

In wild-type cells, AKAP450 and CDK5Rap2 associate with the GA. In agreement with published work [16], we found that removal of AKAP450 dissociated CDK5Rap2 from the GA whereas loss of CDK5Rap2 did not affect Golgi-association of AKAP450, thus confirming that CDK5Rap2 is recruited to the GA through AKAP450 interaction (not shown). Surprisingly, despite the fact that PCNT is not usually detected at the GA, PCNT depletion apparently increased both AKAP450 and CDK5Rap2 labeling at Golgi membranes (Fig. 2A). To quantify this phenotype, we determined the percentage of co-localization between foci of each protein and the Golgi marker GMAP210 in NZ-treated cells (see M&M for details). As shown in Fig. 2B, in the absence of PCNT, association of either AKAP450 or CDK5Rap2 with Golgi membranes enhanced by 1.2 or 1.45 times, respectively.

**Figure 2.**
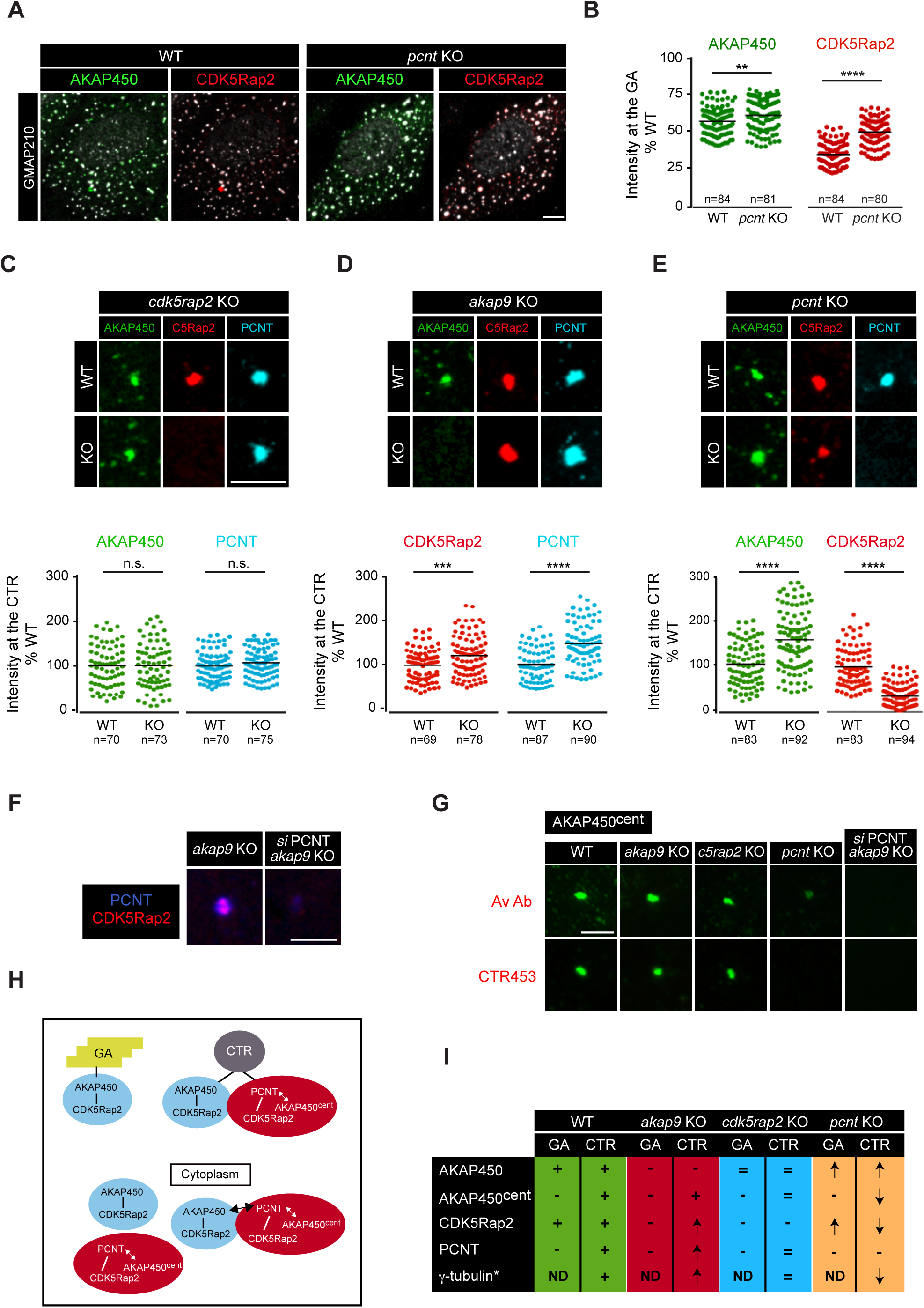
Subcellular distribution of AKAP450, PCNT and CDK5Rap2 in KO cell lines. (**A**) IF images of WT and *pcnt* KO RPE-1 cells treated with NZ for 3 h and triple stained for AKAP450 (green), CDK5Rap2 (red) and GMAP210 as a Golgi marker (white). Double labelings are shown as indicated. (**B**) AKAP450 and CDK5Rap2 fluorescence intensity colocalization with Golgi membranes (GMAP210) in WT and *pcnt* KO cells. Cells were treated and processed as in (A). At least 80 randomly selected cells from each sample were analyzed and the percentages of colocalization determined. Each individual data point represents a cell and the horizontal black lines bars represent the mean. Data were collected from two independent experiments. ** p<0.01; **** p<0.0001 (unpaired two-tailed Student’s t-test). (**C-E**) (Top) High magnification images of centrosomes in WT and either *cdk5rap2* KO (C), *akap9* KO (D) and *pcnt* KO (E) RPE-1 cells labeled for PCM proteins as indicated. C5Rap2: CDK5Rap2. (Bottom) Quantitative analysis of AKAP450 and PCNT (C, n.s.), CDK5Rap2 and PCNT (D, *** p<0.001 **** p<0.0001), or AKAP450 and CDK5Rap2 (E, **** p<0.0001) at the centrosome in WT and KO cell lines as indicated. A region of interest of 1,5 μm radius was drawn around a centrosome marker (CAP350), fluorescence intensities were quantified and normalized to the wild-type control. Data were collected from two independent experiments. (**F**) *akap9* KO RPE-1 cells were transfected with either control (left) or PCNT (right) siRNAs and double stained for CDK5Rap2 (red) and PCNT (blue). In the absence of both AKAP450 and PCNT, no CDK5Rap2 signal was detected at the centrosome. (**G**) WT and all the three KO cell lines were analyzed for the presence of the centrosomal AKAP450 isoform (AKAP450^cent^) with Av and CTR453 antibodies. *akap9* KO RPE-1 cells transfected with PCNT siRNAs were also included in the analysis. (**H**) Schematic representation of proposed AKAP450-based and PCNT-based complexes and their subcellular localization. (**I**) Table summarizing changes observed in centrosome- (CTR) or GA-associated pools of AKAP450, AKAP450^cent^, CDK5Rap2, PCNT and γ-tubulin (*indicates that corresponding data are shown in Fig. 3) in the respective cell lines with respect to WT cells. (+) present, (-) absent, (=) unchanged, (**↑**) increased, (**↓**) decreased, (ND) non-determined. Scale bars= 5 μm.

Regarding the centrosomal levels of AKAP450 or PCNT, no changes were detected in cells lacking CDK5Rap2 demonstrating that this protein is located downstream in PCM association as well (Fig. 2C). On the contrary, knock-out of either AKAP450 or PCNT strongly modified the distribution of the other proteins. Loss of AKAP450 resulted in a 1.5-fold increase of PCNT (Fig. 2D) and reciprocally, depletion of PCNT increased AKAP450 at almost the same degree (Fig. 2E). Although AKAP450 and PCNT have been reported to interact each other [12], these results suggest that they are also able, at least partially, to independently target the centrosome. On the other hand, knock-out of PCNT greatly reduced (70%), but did not eliminate, the binding of CDK5Rap2 to the centrosome (Fig. 2E) confirming that PCNT is an important targeting factor for CDK5Rap2 but not the only one. To test whether AKAP450 also recruits CDK5Rap2 to the centrosome, as it does to the GA, we performed siRNA of PCNT on *akap9* KO cells. As shown in Fig. 2F, CDK5Rap2 was fully displaced from the centrosome when both PCNT and AKAP450 were absent. Finally, we analyzed the behavior of the centrosomal AKAP450^cent^ isoform in the different KO cell lines by using Av and CTR453 antibodies (Fig. 2G). We found that AKAP450^cent^ was retained at the centrosome after removal of AKAP450 and CDK5Rap2 but lost in the absence of PCNT. This suggests that AKAP450^cent^ does not directly bind the centrosome but does it through PCNT interaction.

Taken together, these results support the existence of, at least, two types of γ-tubulin-recruiting protein complexes: those containing AKAP450 and CDK5Rap2 (AKAP450-based complexes) that are present at both the GA and the centrosome and those formed by PCNT, AKAP450^cent^ and CDK5Rap2 (PCNT-based complexes) that specifically localize at the centrosome (Fig. 2H). Significant pools of PCNT-based and AKAP450-based complexes are also present in the cytoplasm from which they can be easily isolated by co-immunoprecipitation (Fig. S3D). Immunoprecipitated complexes contained γ-tubulin (Fig. S3E). However, whether cytoplasmic PCNT-based complexes contained the AKAP450^cent^ isoform could not be determined by WB due to lack of reliable antibodies. Finally, it is possible that all of these proteins form part of the same cytoplasmic complexes through PCNT-AKAP450 interaction. Intriguingly, our findings reveal an inverse relationship between AKAP450-based and PCNT-based complexes at the centrosome: the absence of one of them favors the accumulation of the other one (see Fig. 3F). Although other scenarios are possible, the most plausible explanation for these observations is that PCNT and AKAP450 compete for PACT-domain binding sites at the PCM. A summary of these results is presented in Fig. 2I.

**Figure 3.**
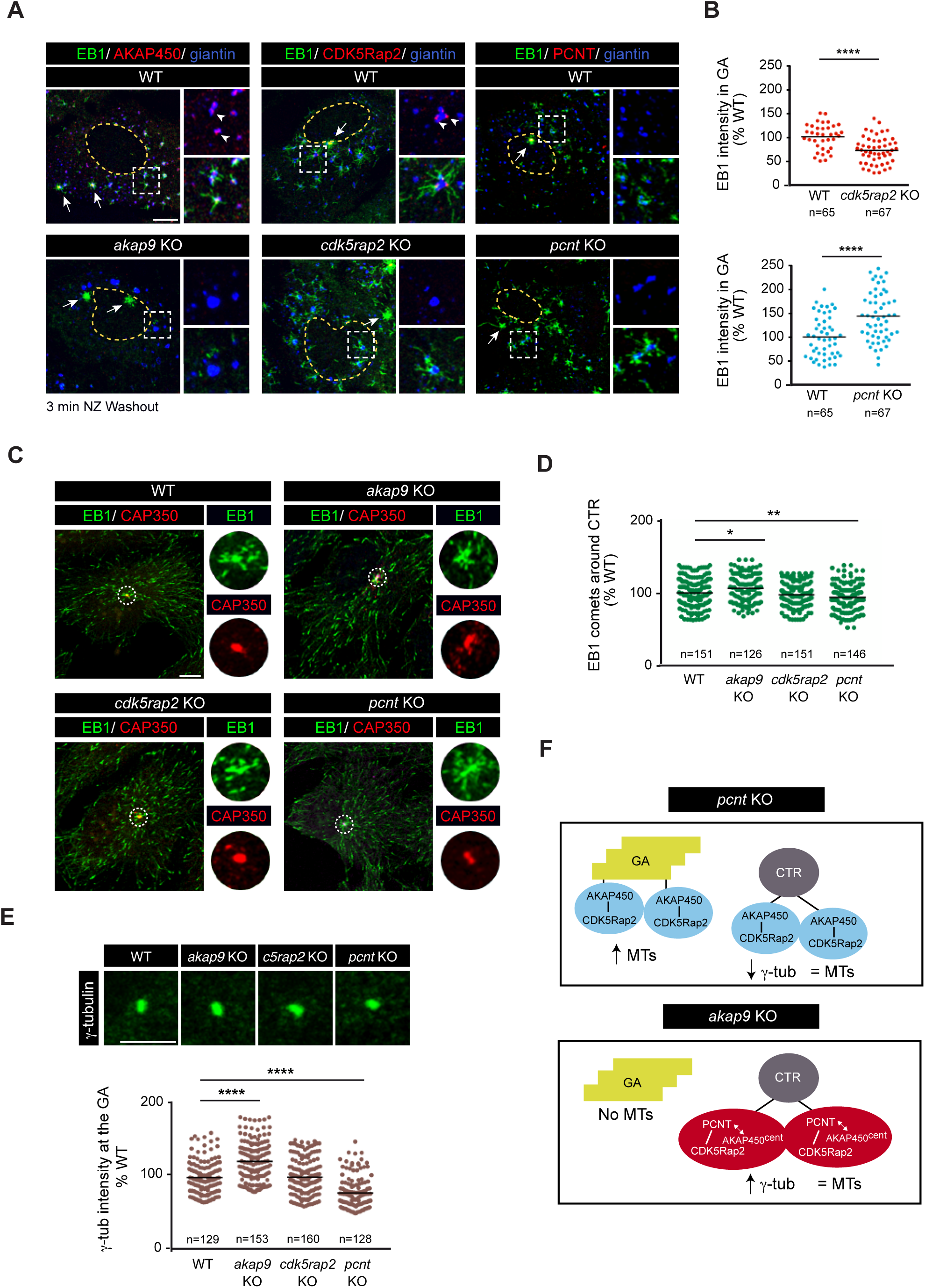
Centrosomal MT-nucleation is independent of the GA. (**A**) MT regrowth experiments in wild-type (WT) and either *akap9* KO (left), *cdk5rap2* KO (middle) or *pcnt* KO (right) cells. Cells were stained as indicated. At right, enlarged images of boxed areas show double stainings for AKAP450/giantin, CDK5Rap2/giantin or PCNT/giantin (top) and triple labelings (bottom). (**B**) (Top) Quantification of EB1 intensity at the Golgi membranes (giantin) in WT and *cdk5rap2* KO cells. After a 3 h of NZ treatment, MTs were allowed to polymerize for 3 min. Data were collected from two independent experiments and normalized to the WT mean. At least 65 cells were analyzed per condition. **** p<0.0001 (unpaired two-tailed Student’s t-test). (Bottom) Same as above but in WT and *pcnt* KO cells. (**C**) Confocal images of WT, *akap9* KO, *cdk5rap2* KO and *pcnt* KO cells stained for EB1 (green) and the centrosome marker CAP350 (red). High magnification single-channel images of selected areas are shown at right. (**D**) Scatter plot shows quantification of EB1 comets at the centrosome. A region of interest of 3 μm radius around the CAP350 signal was used to count the number of comets. Data were collected from two independent experiments and normalized to WT mean. * p<0.05, ** p<0.01 (one-way ANOVA followed by Dunnett’s multiple comparisons test). (**E**) (Top) High magnification images of γ-tubulin at the centrosome in WT and all the three RPE-1 KO cell lines. C5Rap2: CDK5Rap2. (Bottom) Quantitative analysis of γ-tubulin fluorescence intensity. Each individual data point represents a cell and the horizontal black lines bars represent the mean. Data were collected from two independent experiments. **** p<0.0001 (one-way ANOVA followed by Dunnett’s multiple comparisons test). (**F**) Scheme summing up the changes in AKAP450-based and PCNTbased complexes distribution in *akap9* (top) and *pcnt* (bottom) KO RPE-1 cells and their impact in γ-tubulin recruitment and MT nucleation. Scale bars= 7,5 μm.

### Inhibition of MT nucleation at the GA does not interfere with that of the centrosome

We next evaluated the impact of AKAP450, CDK5Rap2 or PCNT depletion on MT nucleation at either the GA or the centrosome. To monitor MT dynamics at the GA we performed MT repolymerization experiments after NZ treatment. Three min after NZ washout, cells were triple stained for the plus-end binding protein EB1, the Golgi marker giantin and either AKAP450, CDK5Rap2 or PCNT. As expected, MTs grew from both the centrosome and Golgi elements in WT cells (Fig. 3A). A close examination showed that MTs arose from Golgi-associated foci that contained both AKAP450 and CDK5Rap2 (Fig 3A, arrowheads). These foci precisely co-localized with minus ends of growing MTs indicating that they represent active MT nucleation sites on Golgi membranes. As previously reported [19], we found that MT nucleation from the GA was inhibited in *akap9 KO* cells and reduced by a 27% in *cdk5rap2* KO cells (Fig. 3B)[21]. Strikingly, removal of PCNT led to 1.5-fold increase in MT nucleation at the GA (Fig. 3B), consistent with a higher association of AKAP450 and CDK5Rap2 with the GA under these conditions (see Fig. 2B). We conclude that MT nucleation at the GA is strictly dependent on AKAP450, facilitated by CDK5Rap2 and negatively regulated by PCNT.

Then, we investigated MT nucleation activity of centrosomes by counting the number of EB1 comets growing from them in their immediate vicinity (Fig. 3C). EB1 comet number somewhat increased in *akap9* KO (106%), remained unchanged in *cdk5rap2* KO cells (101%) and slightly decreased after PCNT depletion (107%; Fig. 3D). These findings suggest that none of these proteins play a critical role in MT nucleation at the centrosome in spite of their reported γ-TuRC binding properties. Centrosomal γ-tublin levels after ablation of *akap9, cdk5rap2* and *pcnt* genes were also determined by measuring γ-tublin fluorescence intensity at the centrosome (Fig. 3E). Interestingly, removal of AKAP450 increased γ-tublin binding to centrosomes (123%) whereas CDK5Rap2 depletion did not affect it (101%) and PCNT KO reduced it by a 21% (Fig 3F). Thus, AKAP450 and PCNT regulate a significant fraction of centrosomal γ-tublin content that is apparently, however, dispensable for MT nucleation. A cartoon summarizing these results is presented in Fig. 3F: in the absence of PCNT, centrosomal γ-tublin diminished in spite of increased AKAP450 at the centrosome under these conditions (Fig. 3F, top); when AKAP450 is lacking, centrosomal γ-tublin increased in parallel with increased PCNT levels (Fig. 3F, bottom).

### Microtubule nucleation activity of the GA negatively correlates with the number of centrosomes

We further investigated the relationship between the centrosome and GA-associated MT nucleation activity by generating cells without centrosomes or with an extra-number of centrosomes. For that purpose, we treated cells with the PLK4 inhibitor centrinone to induce centrosome loss and then, we removed the drug which triggers a wave of centrosome over-duplication [29]. Centrinone-treated cells appeared larger than control cells under the microscope (see also [21]). To better characterize this phenotype, we allowed control and centrinone-treated cells to attach to different size crossbow-shaped micropatterns. We observed that treated cells preferentially adhered to larger micropatterns than control cells (1100 versus 700 μm^2^; Fig. 4A). Analysis by FACS revealed 27% increase in mean cell volume upon 7-days centrinone treatment demonstrating that treated cells were actually bigger than control cells (Fig. 4B). This percentage is probably underestimated since under these conditions almost 25% cells still contained a centriole. To find out whether cell volume increase is due to changes in cell cycle profile, we measured the volume of isolated G1 and G2 centrinone-treated cell subpopulations. Cells with less than 2n or more than 4n DNA content were excluded from the analysis. In both G1 and G2 cells, volume was higher in the absence than in the presence of a centrosome (24.6% or 21.54, respectively). These results unveil an unexpected role of the centrosome in controlling cell size along the cell cycle that deserves further characterization. We also noticed that cells lacking centrosomes displayed an aberrant nuclear morphology although cell ploidy was apparently unaffected (Fig. S4A-B). To minimize the effects of cell volume increase and nuclear perturbations in our results, we carried out all our experiments by treating RPE-1 cells for only 5-6 days, by selecting cells with apparently normal nuclear morphology and by adjusting the data to cell area if required.

**Figure 4.**
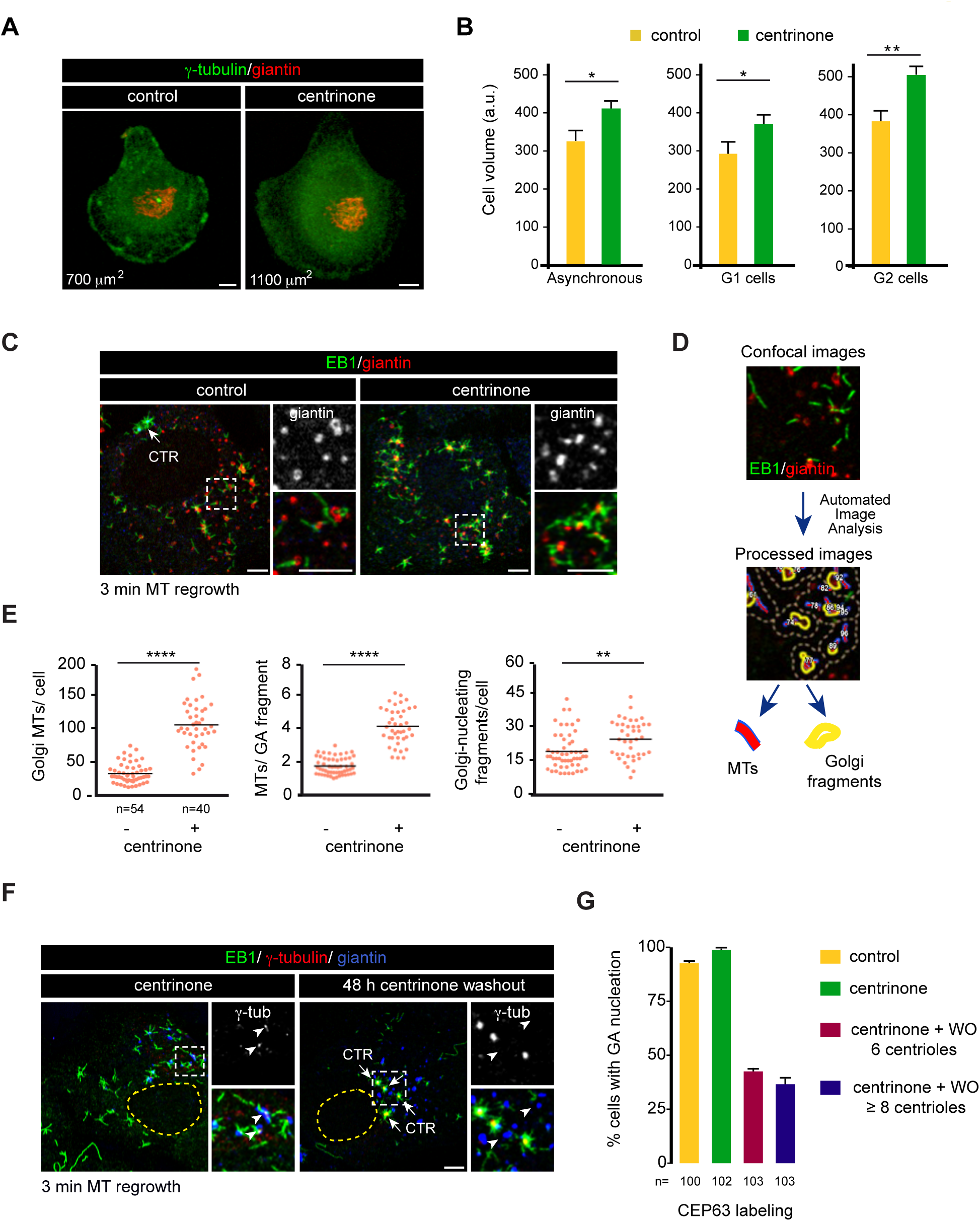
The centrosome regulates the MT-nucleation activity at the GA. (**A**) Examples of control and centrinone-treated cells plated on individual crossbow-shaped micropatterned coverslips and labeled for γ-tubulin (green) and giantin (red). Numbers in μm^2^ represent the approximate cell area depending on the pattern size. (**B**) Quantitative analysis of cell volume by FACS analysis in control and centrinone-treated RPE-1 cells. a.u.: arbitrary units in forward scatter (FSC-H). Bars represent mean values **±** SD of three independent experiments. In the middle and right plots, values correspond to sorted G1 or G2 cells as indicated. * p<0.05; ** p<0.01 (unpaired two-tailed Student’s t-test). (**C**) Control and centrinone-treated cells were subjected to a 3-min MT regrowth assay, fixed and labelled for EB1 (green) and giantin (red). Insets show magnified images of boxed regions. (**D**) Workflow of the method used for quantifying MT nucleation from the GA. (**E**) Quantification of the total number of growing MTs per cell (left), the number of MTs nucleated per Golgi fragment (middle) and the number of MT-nucleating Golgi fragments (right) in both control and centrinone-treated RPE-1 cells. Scatter plots show each individual data point and horizontal black lines bars represent the mean. Data were collected from two independent experiments. ** p<0.01; **** p<0.0001 (unpaired two-tailed Student’s t-test). (**F**) RPE-1 cells were treated with centrinone for either 7 days (left) or for 5 days and then allowed to recover for 48 h in the absence of the drug prior to fixation (right). Representative images of a triple IF staining with EB1, γ-tubulin and giantin as a Golgi marker are shown. Insets show MT nucleation from Golgi elements (left) or from multiple centrioles (right) and are enlarged at right. Arrowheads point to golgi elements. (**G**) Quantification of the percentage of cells exhibiting Golgi-nucleating activity in centrinone-treated cells 48h after the washout of the drug. Cells were divided into two categories depending on the number of centrioles that were visualized by labeling for the centriole protein CEP63. At least 100 cells were analyzed in each condition. The same number of control and centrinone-treated cells were also analyzed for comparison. Scale bars= 5 μm.

When cells lacking centrioles were subjected to MT-regrowth for 3 min after NZ-induced disassembly, virtually all MTs grew as asters from scattered Golgi elements (Fig. 4C). In order to quantitatively assess changes in MT nucleating activity of the GA upon loss of centrioles, we applied a software that allowed automated identification of individual Golgi elements as well as MTs growing from each Golgi element (cartooned in Fig. 4D). Individual cells were delineated before the analysis and centrosomes of control cells were also identified and excluded from the analysis. Quantification of at least 40 control and centrinone-treated cells revealed a three-fold increase in the number of growing MTs/per cell in the absence of centrosome (34.5 MTs/control cell versus 110.5 MTs/centrinone-treated cell). The number of MTs nucleated from each Golgi element increased from 1.8 to 4.3 after centrinone treatment whereas the number of nucleating Golgi elements per cell increased 1.3 times in centrinone-treated cells (20.3 in control versus 26.3 in centrinone-treated cells). This boost in nucleating activity of the GA in the absence of centrosome paralleled the increase in the number of total growing MTs. To rule out that this phenotype could be due to a direct effect of PLK4 inhibition on MT nucleating activity of the GA, we performed similar polymerization experiments in cells treated with centrinone for 3 hours. No differences with control cells were observed (Fig. S4C) confirming that the increase in MT nucleating activity of the GA in centrinone-treated cells is caused by centrosome loss (see also [21]).

We then performed similar experiments 48 h after removal of the drug. Centrinone washout results in the transient hyperactivation of PLK4 that led to the generation of numerous centrosomes [29]. As shown in Fig. 4F, newly formed centrosomes actively recruited γ-tublin and nucleated MTs. On the contrary, the capacity of the GA to bind γ-tublin and to promote MT nucleation was markedly reduced under these conditions. To quantify this phenotype, we determined the percentage of cells containing two (control), none (centrinone-treated), 6 or more than 8 centrioles (centrinone-washout) in which the MT nucleation at the GA was inhibited. New centrioles were identified by Cep63 staining. Fifty-seven per cent of cells containing 6 centrioles did not nucleate MTs from the GA. This inhibitory effect was even stronger in cells containing more than 8 centrioles (63% reduction). From these experiments, we conclude that the MT nucleating capacity of the GA during interphase is not only dependent of the presence of a centrosome but is also affected by the number of centrosomes. Under these conditions, the high rate of MT nucleation at centrosomes apparently dramatically reduces the capacity of Golgi membranes to efficiently assemble MTs.

### Loss of centrioles induces recruitment of PCNT, AKAP450^cent^ and γ-tubulin to the GA

To investigate the factor/s responsible for the enhancement of MT nucleating activity of the GA in the absence of centrosomes, we first examined by IF staining the distribution of AKAP450, CDK5Rap2, PCNT and AKAP450^cent^ in centrinone-treated cells (Fig. 5A). In agreement with early data [29], we observed that the GA exhibited normal morphology and location despite the absence of centrosomes (Fig. 5A). AKAP450 and CDK5Rap2 stainings were also apparently unchanged. However, PCNT and AKAP450^cent^ whose labelings are mostly restricted to the PCM in control cells, appeared as numerous spots concentrated around the GA in cells lacking centrosome (Fig. 5A)

**Figure 5.**
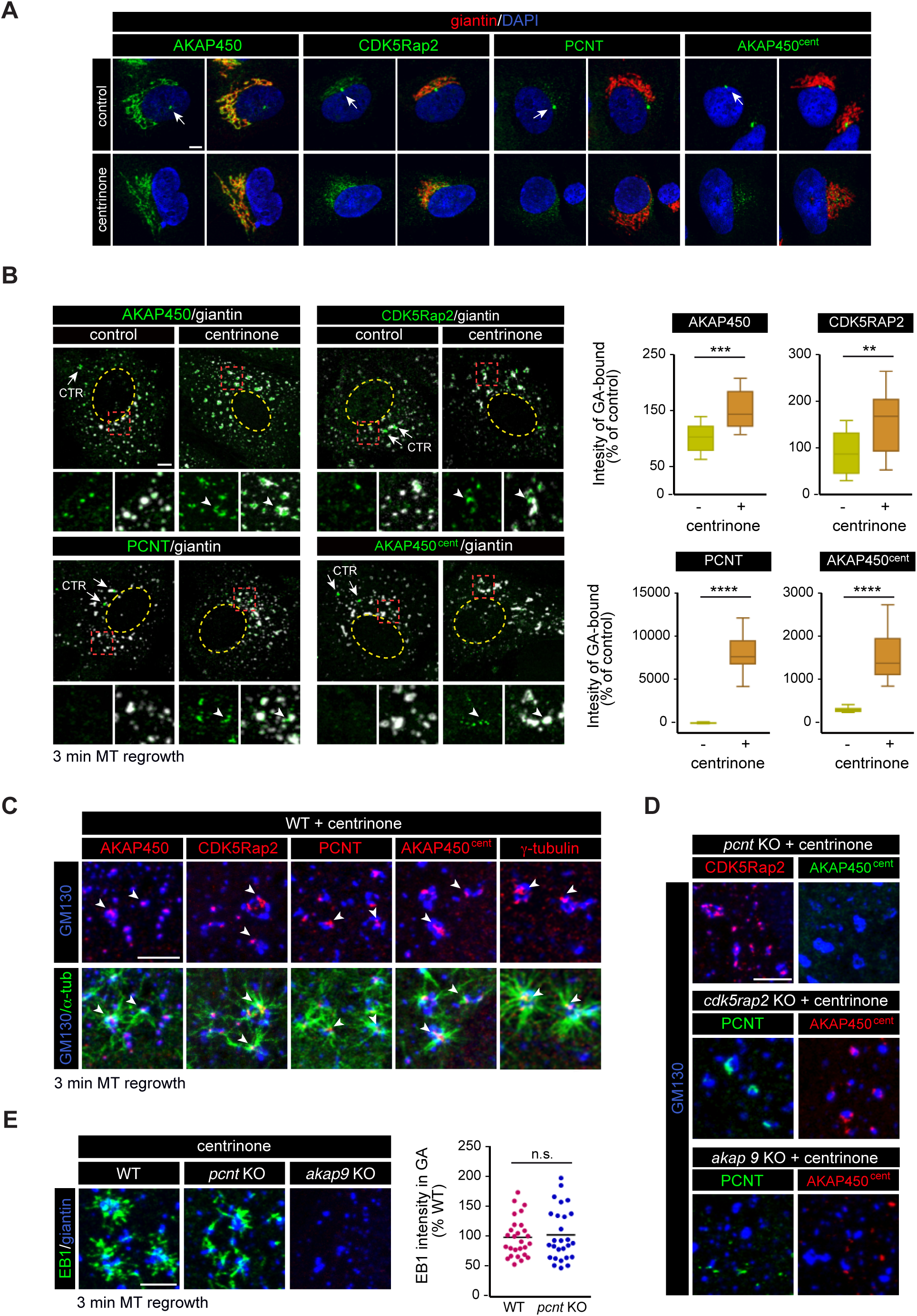
PCM proteins redistributed to the GA in the absence of centrioles. (**A**) Control (top) and centrinone-treated (bottom) RPE-1 cells double stained with either anti-AKAP450, CDK5Rap2, pericentrin or AKAP450^cent^ antibodies (all shown in green) and anti-giantin antibody (red) as a Golgi marker. DNA was counterstained with DAPI and is shown in blue. Single channel (left, green) and merged images (right) are shown in each case. White arrows indicate the centrosome. (**B**) (Left) IF images of control and centrinone-treated RPE-1 cells treated with NZ to induce GA fragmentation and double stained with either anti-AKAP450, CDK5Rap2, PCNT or AKAP450^cent^ (all shown in green) and anti-giantin antibody (white). Enlarged views of the boxed areas are presented at the bottom with or without the signal of the Golgi marker. The yellow dashed line indicates the contour of the nucleus. Arrows indicate the centrosome (CTR) and arrowheads accumulation of the respective proteins in Golgi membrane surfaces. (Right) Box-and-whisker plots showing quantification of association between indicated proteins and Golgi elements (see Materials and Methods for details). Top and bottom ends of the boxes represent 75th and 25th percentiles, and whiskers represent 90th and 10th percentiles. The median is depicted with a solid line. Individual Golgi elements from at least 14 cells were delineated (>500 elements/experiment) and the intensity of each protein spot associated with each Golgi element was measured. Data were collected from two independent experiments and normalized to WT cells. ** p<0.01; *** p<0.001; **** p<0.0001 (unpaired two-tailed Student’s t-test) (**C**) MT-regrowth experiment after NZ treatment in centrinone-treated cells. High magnification images at the bottom are representative of a triple staining with α-tubulin, GM130 and any of the PCM proteins as indicated and show MTs growing from the surface of the GA. Double labelings without α-tubulin are also shown (top) to allow better visualization of PCM-protein spots on Golgi elements acting as MT nucleation sites (arrowheads). (**D**) Centrinone-treated *pcnt* KO, *akap9* KO and *cdk5rap2* KO cells, were incubated with NZ for 3 h, fixed and triple labeled for either CDK5Rap2, AKAP450^cent^ or PCNT as indicated. (**E**) (Left) MT regrowth assay in centrinone-treated WT, *pcnt* KO and *akap9* KO RPE-1 cells stained with antibodies to EB1 and giantin. (Right) Quantification of EB1 intensity at the Golgi membranes in WT and *pcnt* KO cells treated with centrinone, as a measure of MT-nucleation from the GA. The data show no significant differences between the two groups (n.s.). Data were collected from two independent experiments and normalized to the WT control. Scale bars= 5 μm.

A closer examination and quantification after NZ-induced MT disassembly revealed that levels of Golgi-bound AKAP450 and CDK5Rap2 significantly increased in centrinone-treated cells (151% and 154%, respectively; Fig. 5B). In addition, PCNT and AKAP450^cent^ became associated with Golgi elements in centrosome-less cells (81-fold and 15-fold increase, respectively; Fig. 5B). As a control, we also investigated the distribution of the PCM protein Cep192, that has been reported to play a role in centrosomal MT nucleation [31]. We did not observe any accumulation of Cep192 at the GA in centrinone-treated cells (Fig. S5A), indicating that association of PCNT and AKAP450^cent^ to Golgi membranes is not due to redistribution of the whole PCM to the GA in the absence of centrioles, but specifically concerns AKAP450- interacting proteins. MT-regrowth experiments revealed that PCNT and AKAP450^cent^ specifically redistributed to Golgi-associated MT nucleation foci where AKAP450, CDK5Rap2 and γ-tublin accumulated as well (Fig. 5C). These observations support the notion that, in the absence of centrosome, all of these proteins form large MT nucleating complexes at the surface of *cis*-Golgi membranes.

We next wondered how PCNT and AKAP450^cent^, which do not bind Golgi membranes under normal conditions, were addressed to the GA in centrosome-less cells. To answer this question, we analyzed the distribution of these proteins in either *pcnt*, *cdk5rap2* or *akap9* KO cells treated with centrinone. It must be noted that after 7-days of centrinone treatment, *cdk5rap2* and *pcnt* KO cell cultures contained very few cells without centrosome. For unknown reasons, centrinone-treatment of either *pnct* or *cdk5rap2* KO cells had a strong negative impact in cell survival and consequently, less cells could be analyzed in these experiments. In the absence of PCNT, AKAP450^cent^ dissociated from Golgi membranes, as occurs at the centrosome, whereas CDK5Rap2 remained attached (Fig. 5D). Removal of CDK5Rap2 did not alter Golgi membrane association of either PCNT or AKAP450^cent^ (Fig. 5D). Finally, in the absence of AKAP450, all the three proteins dissociated from the GA (Fig. 5D). Altogether these results suggest that in cells lacking centrioles PCNT-based complexes redistribute from the PCM to the GA through interaction of PCNT with AKAP450. Finally, we investigated the consequences of such a redistribution on MT formation from Golgi membranes. Data revealed no significant differences in centrinone-treated *pcnt* KO cells compared with centrinone-treated WT cells (Fig. 5E), indicating that PCNT-based complexes are not required for the enhanced MT nucleating activity of the GA induced by centrinone-treatment.

### In the absence of both centrosome- and Golgi-associated MT nucleation, numerous acentriolar cytoplasmic MTOCs organize the MT network in a PCNT-dependent manner

An unexpected finding was that RPE-1 cells were able to generate a cell-wide MT network when both centrosome and Golgi-mediated MT nucleation were suppressed, i. e. centrinone-treated *akap9* KO cells. MT repolymerization experiments under these conditions revealed asters of growing MTs distributed throughout the cytoplasm that did not co-localize with Golgi membranes. Instead, they arose from numerous and size-variable PCNT-containing cytoplasmic aggregates (Fig. 6A). After 3 h of NZ-washout, MTs organized a dense, non-radial and highly abnormal network Fig. S5B)

**Figure 6.**
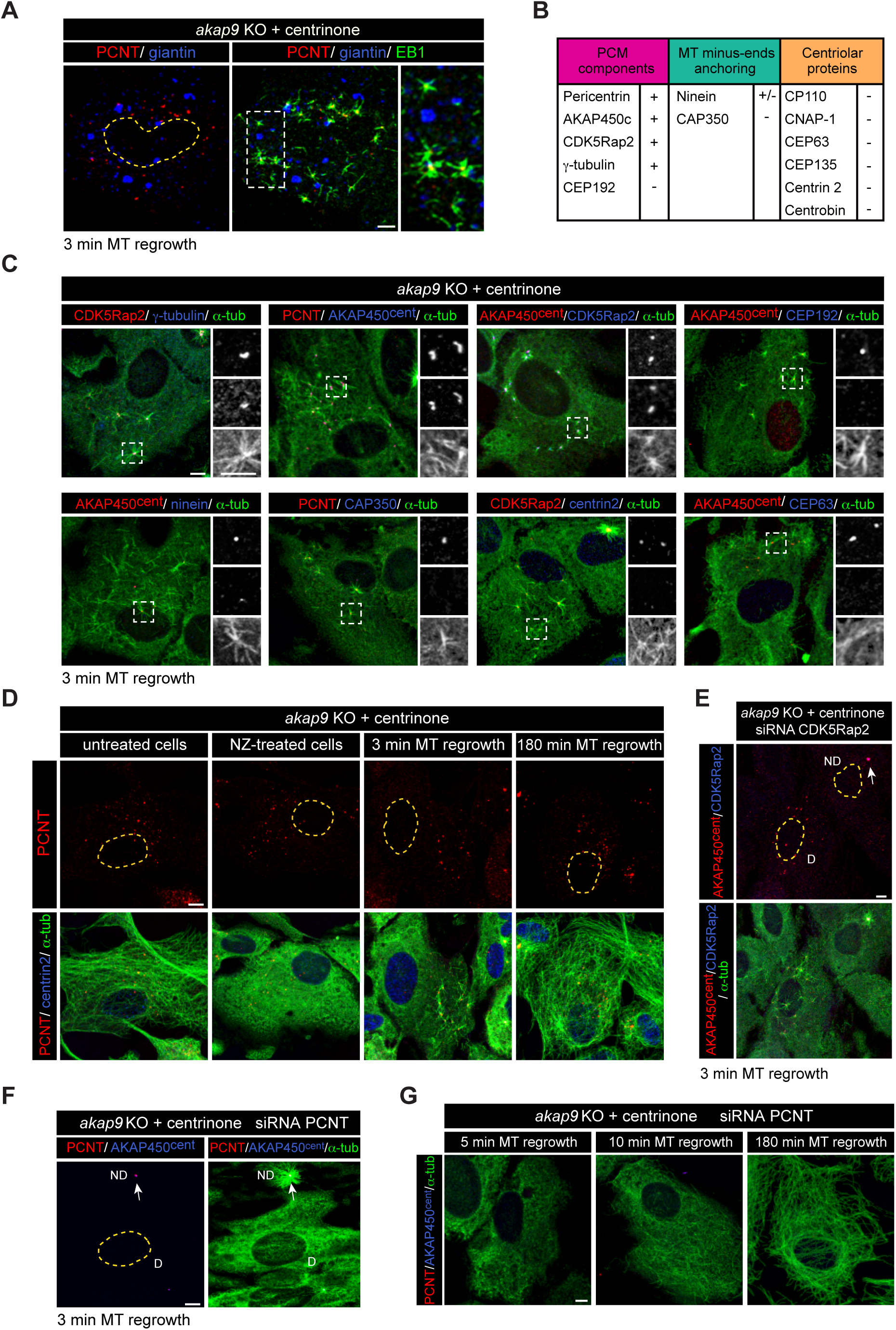
PCNT-dependent acentriolar cytoplasmic MTOCs nucleate MT in the absence of centrosome- and Golgi-mediated MT nucleation. **(A)** MT-regrowth experiment after NZ treatment in centrinone-treated *akap9* KO cells. A representative confocal image of a double labeling for PCNT and giantin is shown in the left panel and the corresponding merged image with EB1 in the middle panel with enlarged view at right. Note that MT asters growing in the cytoplasm are not associated to Golgi fragments. The dashed line in the left panel corresponds to the contour of the nucleus **(B)** Summary table of the proteins present (+) or excluded (-) from the cytoplasmic aggregates. **(C)** Centrinone-treated *akap9* KO cells were subjected to a MT regrowth assay and triple stained with the indicated antibodies. High magnification images of the boxed areas are shown as individual labelings in black and white. Top panels show the red marker, middle panel the blue marker and bottom panel the green marker. **(D)** MTs were depolymerized by NZ treatment and, at the indicated time points after washout, cells were fixed and labeled for α-tubulin (green), PCNT (red) and centrin 2 (blue). Single labelings for PCNT are shown on the top and the corresponding merged images at the bottom. **(E, F)** *akap9* KO cells were treated with centrinone in order to induce cytoplasmic aggregates and then transfected with siRNAs specific for CDK5Rap2 (E) or PCNT (F). Cells were further subjected to a 3-min MT regrowth assay. In (E) cells were stained for AKAP450^cent^ (red), CDK5Rap2 (blue) and α-tubulin (green). In (F) cells were labeled for PCNT (red), AKAP450^cent^ (blue) and α-tubulin (green). D indicates a depleted cell. ND indicates a non-depleted cell that still contains a centriole. **(G)** MT repolymerization experiments of centrinone-treated *akap9* KO cells transfected with RNA duplexes specific for PCNT. At indicated time points, cells were fixed and labeled for PCNT (red), AKAP450^cent^ (blue) and α-tubulin (green). Yellow dashed lines indicate the nuclei contours. Scale bars= 5 μm.

To further characterize these cytoplasmic MTOCs, we tested a panel of antibodies against PCM proteins, minus-end anchoring proteins and centriole associated proteins (resumed in Fig. 6B). As shown in Fig. 6C, aggregates contained PCNT, AKAP450^cent^, CDK5Rap2 and γ-tublin but lacked Cep192. The MT anchoring protein CAP350 was also absent while the presence of ninein was variable depending on the size of the aggregate, being visible only in some of the larger ones (Fig. 6C and Fig S5C). Notably, asters growing from these bigger aggregates were more focused than the rest (Fig. S5C). None of the centriolar markers tested, i.e., CEP63, centrin-2, (Fig. 6C), CEP135, CP110, CNAP-1 and centrobin (Fig. S5C) were detected in the aggregates. These results suggest that cytoplasmic aggregates are similar to centrosomal PCNT-based complexes and behave as true acentriolar cytoplasmic MTOCs. They also demonstrate that they are fully competent to nucleate and anchor MTs and, in this way, to generate a cell-wide array of MTs.

We next monitored the dynamics, if any, of the acentriolar MTOCs during our NZ-washout experiments (Fig. 6D). We did not observe any variation neither in the number nor in the size of acentriolar MTOCs that remained scattered throughout the cytoplasm all along the MT polymerization process. Thus, cytoplasmic aggregates exist both in the absence of MTs and in fully assembled MT networks indicating that they represent stable MT-nucleating modules.

Finally, in an attempt to identify factors required for the formation of acentriolar MTOCs, we performed either CDK5Rap2 or PCNT siRNA experiments on *akap9* KO cells that had been treated with centrinone (Fig. 6E and F). Once again, knocking-down CDK5Rap2 did not prevent either the formation of acentriolar cytoplasmic MTOCs nor MT nucleation from them (Fig. 6E). In contrast, in the absence of PCNT neither cytoplasmic MTOCs nor asters of growing MTs could be detected three minutes after NZ washout (Fig. 6F). Since CDK5Rap2 was not strictly required for efficient MT nucleation from cytoplasmic aggregates, the minimal MT-nucleating module identified in this work consists of PCNT, AKAP450^cent^ and g-tubulin. Interestingly, depletion of PCNT on centrinone-treated *akap9* KO cells delayed, but did not block, formation of MTs. After ten minutes of NZ washout scattered individual MTs were visible throughout the cytoplasm and, by 3h a disorganized meshwork of MTs completely filled the cytoplasm (Fig. 6G).

### Centrosomes negatively regulates MTOC activity of other subcellular sites

Finally, we wanted to evaluate the impact of manipulating MT nucleation process on the overall MT network organization. To do so, we labeled WT, *akap9*, *cdk5rap2* and *pcnt* KO cells, treated or not with centrinone, for either a-tubulin or EB1 (Fig. 7A-B). On the right of each panel, a cartoon represents active MT nucleation sites under each condition tested. Individual cells were delineated and cell area, MT mass polymer and MT growth were quantified. MT mass was estimated by quantifying total a-tubulin fluorescence signal of individual cells under each condition. Data obtained were then normalized with respect to the WT a-tubulin mean signal. Since cells without centrosome were found to be larger than control cells (see Fig. 7C), we also measured cell area under each condition and calculated MT density by referring total MT mass to cell surface. In an attempt to evaluate MT dynamics, we also determined the number of EB1 comets per cell under all conditions.

**Figure 7.**
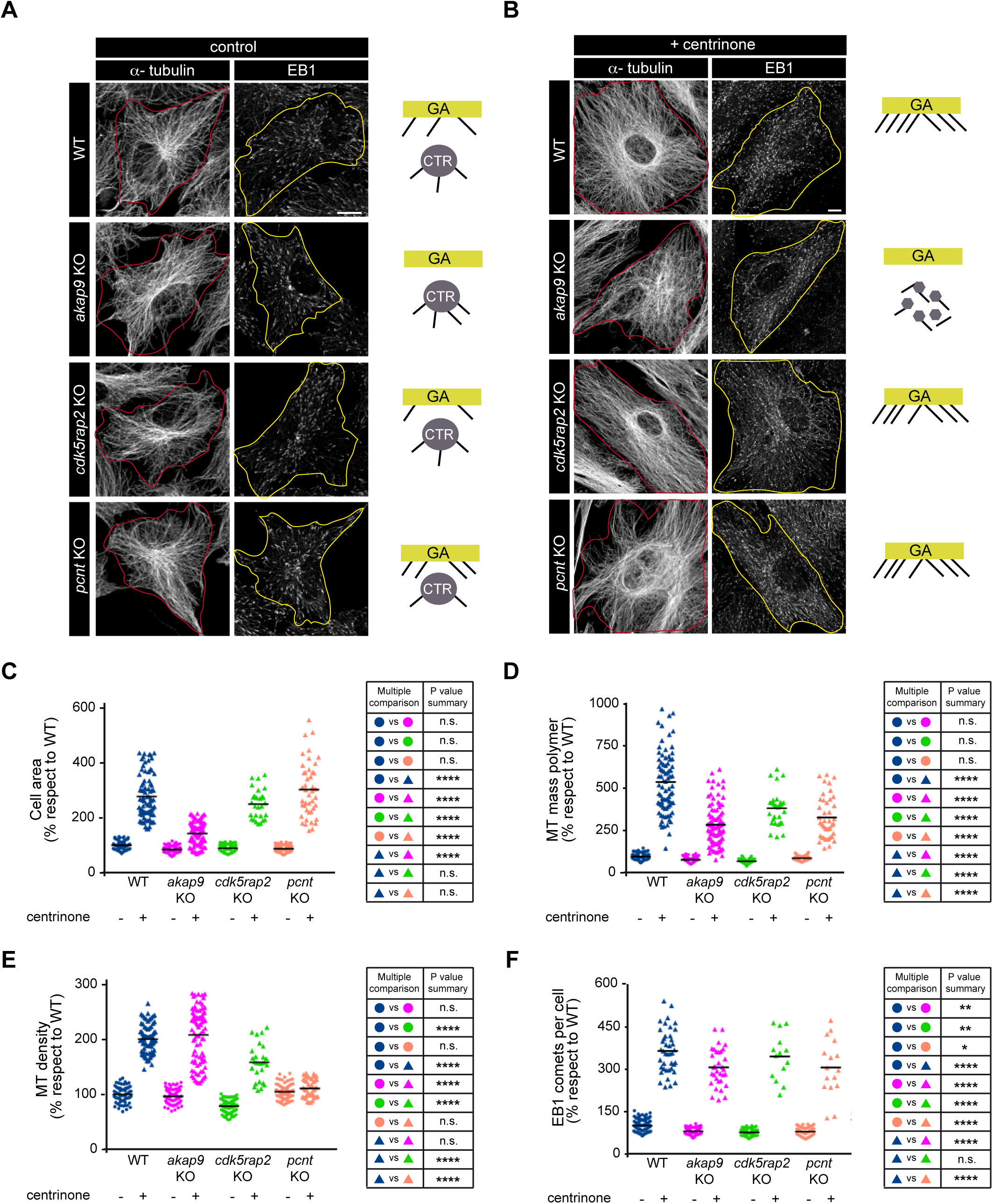
The centrosome negatively regulates MTOC activity of other subcellular sites. (**A**) Confocal images showing MT network in WT, *akap9* KO, *cdk5rap2* KO and *pcnt* KO RPE-1 cells fixed and stained with antibodies against α-tubulin (left) or EB1 (right). The red and yellow lines indicate the contour of the cell. At far right, schematic representation of MT nucleation in each case. GA= Golgi apparatus; CTR= centrosome. (**B**) Same as in (A), but in centrinonetreated cells. (**C**) (Left) Quantification of cell area in control and centrinone-treated WT, *akap9* KO, *cdk5rap2* KO and *pcnt* KO RPE-1 cells. Scatter plot shows each individual data point and horizontal black lines bars represent the mean. Data were collected from three independent experiments and normalized to the untreated WT control. (Right) Summary table showing the most relevant data about statistical significance in the multiple comparisons test. n.s.: non-significant; * p< 0,05; ** p<0.01; *** p<0.001; **** p<0.0001 (One-way ANOVA followed by Tukey’s multiple comparison test). (**D**) Quantitative analysis of α-tubulin fluorescence intensity as a measure of the total MT mass polymer in the same cells as in (C). (**E**) Scatter plot showing quantification of MT density (MT mass polymer/ cell area). Data are presented as above. (**F**) Graph shows quantification of the total number of EB1 comets per cell in control and centrinone-treated WT, *akap9* KO, *cdk5rap2* KO and *pcnt* KO RPE-1 cells. Scale bars= 10 μm.

Depletion of either AKAP450, CDK5Rap2 or PCNT led to a small reduction of the cell area (15%, 10% and 12%, respectively; Fig. 7C). Conversely, loss of centrosome resulted in a 3-fold increase in the occupied surface with respect to non-treated WT cells (276%). This increase is significantly higher than that of cell volume determined by FACS analysis of centrosome-less RPE-1 cells (see figure 4B) suggesting that adhesion to substrate is also probably altered in centrosome-less cells. As a matter of fact, in the absence of centrosome, we observed a similar increase in cell surface for CDK5Rap2-depleted cells (248%), and even more for PCNT-depleted cells (299,8%). Centrinone-treated *akap9* KO cells were also larger than *akap9* KO cells containing centrosome but to a lesser extent (142%). Although additional work is required to further characterize this effect, these results strongly suggest that the centrosome control not only cell volume but also cell adhesion.

Notably, changes in MT polymer mass essentially match those of cell area (Fig. 7D). AKAP450- and PCNT-depleted cells contained 18% or 9% less MTs that WT cells, respectively. Since KO cells are smaller that WT cells, they eventually showed similar MT density to WT cells (96.6% and 104%, respectively; Fig. 7E). These results are consistent with the existence of compensatory mechanisms that contribute to maintain the total number of MTs, as proposed above. More pronounced was the effect of CDK5Rap2 removal on MT density (79.1% of WT) possibly due to its proposed role in MT stabilization (Fig. 7D and E).

Nevertheless, these differences appear almost negligible when compared with the effect of centrosome loss on MT network. Indeed, in the absence of centrosome, MT polymer mass increased by five times and MT density by two times (Fig. 7D and E). Centrinone-treated *akap9* KO, *cdk5rap2* KO and *pcnt* KO cells also contained much higher MT mass than centrosome-containing KO cells (294.1%, 392.8 and 336.8%, respectively; Fig. 7D). MT density values also increased in centrinone-treated cells with respect to non-treated cells except in *pcnt* KO cell line. In that case, the huge increase in cell area buffered that of MT mass, resulting in unsignificant differences in MT density between treated and non-treated cells. Finally, we determined the number of EB1 comets that augmented more than 3 times in the absence of centrosome in both WT and KO cell lines. Taken together, these data strongly support the notion that, independently of where MTs are being nucleated from, the centrosome restricts global MT nucleation activity. Since in WT, *cdk5rap2* and *pcnt* KO cells lacking centrosome, MTs are nucleated from the GA, whereas they grow from Golgi-unbound cytoplasmic acentriolar MTOCs in centrinone-treated *akap*9 KO cells, we conclude that the centrosome is able to limit MT nucleating activity on any subcellular structure. From this data, the possibility also arises that PCNT play a role in the inhibitory pathway connecting the centrosome with the GA.

## DISCUSSION

The possibility to easily obtain cells without centrosome, using the reversible drug centrinone [29] and to generate PCNT-, AKAP450- and CDK5Rap2-depleted cells has enabled us to investigate global MT assembly regulation. We found that centrosome activity, which can be modulated by varying the number of centrosomes, regulates the rate of GA-associated MT nucleation, and that both nucleating organelles inhibit MT nucleation from cytoplasmic MTOCs. We conclude that the centrosome determines the total amount of cellular MTs, as centrosome-less cells contain many more MTs than control cells. This unveils an unexpected role of the centrosome as an inhibitor of MT nucleation elsewhere in the cell.

### Regulation of MT nucleation during interphase

Our results reveal substantial differences between MT nucleation regulation at the centrosome and the GA. Neither PCNT, nor AKAP450 or CDK5Rap2 proteins were required for MT nucleation at the centrosome while MT nucleation at the GA fully depended on AKAP450. Depleting CDK5Rap2 only modestly reduced MT assembly at the GA, suggesting that CDK5Rap2, like its paralog myomegalin [21], might contribute to other aspects of MT formation such as stabilization or anchoring. That MT nucleation capacity of the GA was stimulated by either PCNT depletion or centrosome loss, whereas it was inhibited by a high centrosome number, is however intriguing. Furthermore, MT density increase induced by centrosome-loss in WT, AKAP450 or CDK5Rap2-depleted cells did not take place in PCNT lacking cells. Although far from being understood at the mechanistic level, our results point to PCNT as a possible important actor in the regulatory role exerted by the centrosome on GA MT nucleating activity.

The scenario appears even more complex at the centrosome where PCNT-based complexes recruited γ-TuRCs whereas AKAP450-based complexes did not. Furthermore, recruitment of centrosomal PCNT and AKAP450 inversely correlate, probably by competing for PACT domain binding sites at the PCM. We estimate that up to 40% of centrosomal γ-tublin depends on the balance between PCNT and AKAP450, the latter protein playing an inhibitory role on γ-tublinrecruitment. Strikingly, neither depleting nor increasing centrosomal PCNT did alter MT nucleation suggesting that γ-TuRCs bound to PCNT-based complexes are inactive at the centrosome. Similarly, CDK5Rap2 depletion had no effect on γ-tublin recruitment or MT nucleation rate at the centrosome. These results contrast with reports claiming that PCNT, and specially CDK5Rap2, recruit γ-TuRCs and promote MT nucleation [14, 32]. A possible explanation for such discrepancies is that these proteins play different roles depending on the organism, the cell cycle stage or the cell type [28, 33, 34].

Nevertheless, a more systematic analysis is required to know the real contribution to these γ-TurC binding proteins to centrosomal function in different situations. In sum, while MT nucleation at the GA fully relies on AKAP450-based complexes, the contribution of PCNT- and AKAP450-based complexes to MT nucleation at the human centrosome during interphase seems to be minor, or even negligible. What are then the functions of these complexes at the centrosome? It is tempting to speculate that, in addition to their documented roles on organizing the PCM and in centrosome maturation [35], they enable centrosome to somehow sense MT nucleation at other sites due to the capacity of both AKAP450 and PCNT to act as signaling platforms [36]. This work has also identified AKAP450^cent^, a strictly centrosomal isoform of AKAP450, whose exact function at the centrosome will have to be worked out.

### Hierarchical regulation of MT nucleation in mammalian cells

This study reveals a hierarchy in the regulation of the MT nucleation process in RPE1 cells during interphase. Loss of centrioles doubled MT nucleation activity of the GA. Under these conditions, MTs grew from discrete Golgi-associated foci that contained higher number of AKAP450-based complexes and γ-tubulin. Strikingly, centrosomal PCNT-based complexes also redistributed to these Golgi-associated foci in centrinone-treated cells although they did not apparently contribute to the increased MT nucleation activity of the GA. These finding suggests that recruitment of PCNT-based complexes to the GA is normally restricted by centrioles and that these complexes are inactive at both subcellular locations. On the contrary, when both centrosomes and AKAP450 are lacking, cytoplasmic PCNT-based complexes acquire MT nucleating ability, thus acting as cytoplasmic acentriolar MTOCs.

These acentriolar cytoplasmic MTOCs were able to organize a cell-wide MT network suggesting that they contain all the activities required for the successful formation of MTs. However, among all centrosomal markers tested, we only detected PCNT, the AKAP450^cent^ isoform described here, CDK5Rap2, γ-tublin and, sporadically ninein, that could interact with ***γ***-tubulin-containing complexes or with PCNT itself [37, 38]. Cytoplasmic MTOCs did not contain Cep192, a PCNT-interacting scaffolding protein reported to be important for γ-tublin recruitment at the centrosome [39]. Cytoplasmic MT nucleation foci are known to be present in early blastomeres of mouse embryos which lack centrioles during the first embryonic divisions [40] and in centrosome-less cells depleted of the ubiquitin ligase TRIM37 [41]. In the latter case, however, the nucleating foci contained Cep192 and Cep152 but lacked y-tubulin, PCNT and CDK5Rap2. They are, therefore, essentially different from acentriolar MTOCs reported in this work (see also [42, 43]). A linear acentriolar structure was also reported to develop in cells lacking centrioles, AKAP450 and CAMSAP2, that was able to nucleate a dense MT network [21]. In any case, these observations unveil the ability of PCM proteins to assemble in different kinds of MT-nucleating complexes when centrioles are absent. Notably, when all MT nucleation is apparently inhibited, i.e., cells lacking centrosomes, AKAP450 and PCNT, numerous individual MTs formed throughout the cytoplasm that eventually organized a MT network. This agrees with recent data showing that inhibition of γ-tublin by gatastatin does not affect MT density; it decreased MT dynamics but not the number of growing MTs [44]. It is possible that spontaneously assembled MT seeds became stabilized by cytoplasmic factors such as CAMSAP proteins, at the minus ends [45], and +TIPs proteins (CLASPs, CLIP150, p150*^glued^*, etc) at the plus ends [46].

Somewhat paradoxically, cytoplasmic MTOCs are not generally observed in control cells, in spite of the abundant cytoplasmic pools of AKAP450, PCNT, CDK5Rap2 and y-tubulin. Neither did they appear in centrinone-treated cells. As mentioned above, targeting of γ-TuRCs to MTOCs is considered to be crucial for MT nucleation. Thus, it is generally assumed that γ-TuRC-binding protein-complexes become competent for nucleation only once assembled at the PCM. A similar scenario was proposed for the GA where GM130-mediated targeting of AKAP450- based complexes could restrict the assembly of active MT nucleating complexes to these membranes [19]. However, the ability of the cytoplasmic acentriolar MTOCs to nucleate MTs and to generate a cell-wide MT network suggest that targeting to a subcellular structure might not be strictly required. In an alternative and speculative model, activity of cytoplasmic PCNT-and AKAP450-based complexes would be actively suppressed by the presence of inhibitory components, or by lack of activators. Centriole-mediated suppression of MT nucleation in the cytosol has been theoretically predicted [47], in the light of observations on mitosis of early blastomeres from *C. elegans*. Our data during interphase agrees with this idea, providing MT nucleation activity at the GA is also inhibited. In that case, both centrioles and Golgi membranes function as inhibitors of cytosolic MT nucleation.

### MT nucleation and cell differentiation

It is known that the MT nucleating activity of the centrosome is down-regulated during differentiation of epithelia, neurons or pancreatic cells [25-28], or even that centrosomes can be eliminated like in muscle mammalian cells [48]. In most cases, MT assembly is then ensured by Golgi elements [5]. Although molecular mechanisms underlying the loss of MT nucleating activity at the centrosome during cell differentiation remain largely unknown, a common feature to several cell types is the shedding of PCM components, including PCNT, CDK5Rap2 and γ-tubulin. An attractive hypothesis would be that shedding of PCM proteins from centriole backbone during differentiation results not only in inactivation of the centrosome, but also in activation of nucleation at the GA as it occurs in centrinone-treated cells. Interestingly, Golgi localization of PCNT has been reported in terminally differentiated skeletal muscle fibers [26] where MT nucleation occurs at both the nuclear envelope and the GA. Skeletal muscle cells have lost their centrosomes [48], a physiological situation that is mimicked by centrinone-treated cells. The possibility that GA-associated MT nucleation might also be down-regulated, leading to the formation of cytoplasmic acentriolar MTOCs, is so far a mere speculation. But this would confer plasticity to the MT nucleation process and thus would facilitate the re-organization of complex MT arrays in terminally differentiated cells. At any rate, mechanisms unveiled in the present work are likely to be relevant for understanding MT network reorganization during cell differentiation.

### Towards a reappraisal of the role of the centrosome organelle in animal cells

As discussed above, MT nucleation appears as a hierarchically controlled process with the centrosome upstream of the regulatory cascade. Suppression of the centrosome strongly stimulated Golgi MT nucleation whereas inhibition of GA-associated MT nucleation did not affect nucleating activity of the centrosome. Remarkably, amplification of centrosome number inhibited MT nucleation at the GA. Downstream of the cascade, cytoplasmic MT nucleation required blocking both centrosome- and GA nucleating activities.

Particularly surprising was the finding that both centrosome-less wild-type cells, in which the GA becomes the only MTOC, and centrinone-treated *akap9* KO cells, in which MTs are nucleated from cytoplasmic foci, displayed twice higher MT density than control cells. Therefore, centrosome activity seems not only able of controlling the cellular distribution of MT nucleating activity, but also of maintaining a steady state number of MTs in cells. This could be achieved either by down-regulating MT nucleation capacity of alternative MTOCs or by controlling its own activity. These results suggest that the specific organization of the centrosome, in which the centriole barrels control the organization of the PCM, is important and goes beyond the notion of simply concentrating γ-tubulin-containing nucleating complexes. Deciphering the mechanisms which could explain this unexpected role of the centrosome will require a more detailed analysis of the kinetics of nucleation in all conditions described in this work (in progress). Our data adds a new function to the centrosome namely that of governing the total amount of assembled tubulin in cells. Interestingly, this function apparently strongly impacts cellular architecture: we did not observe significant perturbations of cell morphology in the absence of centrioles but cell volume, nuclear morphology and cell area were strongly modified. Increase in MT number and cell volume correlated with a significant increase in cell spreading in cells lacking centrosome. These results suggest that adhesion to substrate is probably dependent on the amount of MTs present in cells, a possibility which fits with the established notion that MTs regulate focal adhesion dynamics [49].

How cells sense their own shape and size is an interesting question that remains poorly understood. Depending on the cellular models, the emphasis is placed either on actin-based dynamic structures in somatic cells [50] or on MT network in eggs [51]. Overwhelming evidence for an integrated mutual regulation of actin cytoskeleton and MTs exist at several levels, including the centrosome [50]. In addition, many kinases, phosphatases and other signaling components, are known to associate with the centrosome, leading to the proposal that centrosomes could act as signaling centers [36]. Ablating the centrosome may thus have direct effects on diverse aspects of cell activity, far beyond those that would be due to impairment of MT nucleation by structural templating.

## MATERIAL AND METHODS

### Cell culture, antibodies and treatments

Immortalized human pigment epithelial cells hTERT-RPE1 (Clontech) and HeLa cells were grown in DMEM/F12 or DMEM, respectively supplemented with 10% FBS at 37°C in 5% CO2. hTERT-RPE1 FRT/TO cells were provided by J Pines (Gurdon Institute, Cambridge, UK). hTERT-RPE1 cells were treated with 125 nM centrinone (kindly provided by K Oegema and A Siau, Ludwig Institute for Cancer Research, La Jolla, CA), for 4-7 days to induce centrosome depletion as previously described [29]. For washout experiments, we treated the cells with centrinone for 5 days and after washout of the drug, cells were maintained for 48 h prior to analysis.

For CYTOO-Chips experiments, control and centrinone-treated cells were seeded on Starter’s CYTOOchips (CYTOO SA, Grenoble, France) following the manufacturer’s protocol. Briefly, trypsinized cells were diluted to a concentration of 12,500 cells/ml, 50,000 cells dispensed into each micropattern and allowed to sediment for 10 min under the hood before moving them to the incubator. After 45 min, the medium was changed, the coverslip surface was gently flushed, and finally, cells were allowed to spread for 5 h before fixation.

SC3-1 and SC3-2 anti-CDK5Rap2 rabbit polyclonal antibodies were generated by Biomedal S.L. (Seville, Spain). To this purpose, constructs containing amino acids 650-900 or 1400-1600 of human CDK5Rap2 were inserted into the expression vector pMAB36-6xHis-LYTAG and the resulting fusion proteins were purified by using the C-LYTAG fusion protein purification system. Rabbit polyclonal anti-AKAP450 (A24), human sera against GMAP210, CTR453 mouse monoclonal antibody, and mouse monoclonal anti-CAP350 have been previously described [19, 52]. CTR453 monoclonal antibody recognizes exon 29 of human AKAP450 [30]. Monoclonal anti-a-tubulin and anti-g-tubulin (clone GTU88) were from Sigma-Aldrich; mouse monoclonal anti centrin-2 (clone 20H5), rabbit polyclonal anti-Cep63 and rabbit polyclonal anti-CDK5Rap2 (Mi) were from Millipore. Mouse monoclonal anti-EB1 and anti-AKAP450 (7/AK) were from BD Biosciences. Mouse monoclonal anti C-Nap1 was from Santa Cruz Biotechnology. Rabbit polyclonal anti-GFP was from ICL, rabbit polyclonal anti-CP110 was from Proteintech Group and rabbit polyclonal anti AKAP9 (Av) was from Aviva Systems Biology. Rabbit polyclonal anti-cep135, mouse monoclonal and rabbit polyclonal anti-PCNT were from Abcam. Human anti-α-tubulin (F2C-hFc2) and human anti-giantin (TA10 hFc2) were from the recombinant antibody platform of the Institut Curie. Rabbit anti-Cep192, anti-GM130 and goat anti-CAP350 were generous gifts from L. Pelletier (Lunenfeld-Tanenbaum Research Institute, Toronto, Canada), Y. Misumi (Fukuoka University, Japan) and E Nigg (Biozentrum, University of Basel, Swizerland), respectively. All secondary antibodies were from Jackson ImmunoResearch.

### Constructions, transfections and RNA interference

The sequences of the stealth siRNAs used were as follows: PCNTB-1: 5′ AAAAGCTCTGATTTATCAAAAGAAG; PCNTB-2: 5′ TGATTGGACGTCATCCAATGAGAAA; AKAP450: 5′ AGTAATTGTTGTTCAACTGGGCCTG; CDK5Rap2: 5′ CCTAAAGCTCCGCATCTATTT. Scrambled siRNA was used as control. Duplexes were obtained from Life Technologies. Assays were performed 72-96h after transfection. GFP-AK1, GFP-AK2, GFP-AK3 and GFP-AK4 fusion proteins have been previously described [20].

Both DNA and siRNA transfections were performed with Neon Transfection System (Invitrogen) by following instructions from the supplier.

### CRISPR/Cas9 mutagenesis of *akap9* and *cdk5rap2* genes in hTERT-RPE1 cells

In order to generate RPE-1 cell lines lacking either AKAP450, CDK5Rap2 or PCNT proteins we first generated a new plasmid based on the Cas9-nickase containing plasmid pSpCas9n(BB)-2A-GFP (PX461) (a gift from Feng Zhang, Addgene plasmid # 48140). We introduced in the XbaI site of PX461 a 409 bpDNA fragment containing a hU6 promoter, sgRNA BsaI cloning sites and a scaffold RNA coding sequence. This fragment was obtained by PCR of pGL3-U6-sgRNA-PGK-puromycin (a gift from Xingxu Huang, Addgene plasmid # 51133) using the primers pFA2 (TCTAGAGAGCGGCCGCCCCCTTCACC) and pFA3 (TCTAGAGTCTCGAGGTCGAGGATTC). The new plasmid named plFA1 contains the features of PX461, an additional cloning site for an sgRNA coding sequence flanked by a hU6 promoter and a scaffold RNA coding sequence. plFA1 allows the expression of the Cas9n protein and two sgRNAs from a single plasmid, thus increasing the efficiency and the specificity of mutagenesis. Primers for targeting the exon-2 of *akap9*, the exon-1 of *cdk5rap2* and the exon-5 of *pcnt* were designed and cloned into plFA1 following Feng Zhang Crispr web tools and protocols (http://crispr.mit.edu/). To facilitate the cloning of complementary primer pairs into plFA1, they were designed with overhand ends homologous to the overhands DNA strands generated by the digestion of plFA1 with BbsI or BsaI enzymes (i. e.). The primer pairs used to generate the sgRNA coding sequences targeting *akap9* exon-2 were CACCGAGAAATAACCAGTCATGAGC/AAACGCTCATGACTGGTTATTTCTC (to be cloned into the BbsI sites of plFA1) and CCGGGTTCTCATTATTGTAGATTC/AAACGAATCTACAATAATGAGAAC (to be cloned into the BsaI sites of pLFA1). The sequences of the primer pairs used to generate the sgRNA coding sequences targeting CDK5RAP2 exon-1 were CACCGAAGAGGACGTCACCGTCCCT/AAACAGGGACGGTGACGTCCTCTTC (to be cloned into the BbsI sites of plFA1) and CCGGCAAGTCCATCATGGCTACAG/AAACCTGTAGCCATGATGGACTTG (to be cloned into the the BsaI sites of pLFA1). The primer pairs used to generate the sgRNA coding sequences targeting *pcnt* exon-5 were CACCGCAACATGCACACGGCGCAGC/AAACGCTGCGCCGTGTGCATGTTGC (BbsI sites) and CCGGGGCGCAGCGCCTCCAGCTCA/AAACTGAGCTGGAGGCGCTGCGCC (BsaI sites). The final plasmids containing the two sgRNAs coding sequences for targeting each of the genes were electroporated in RPE-1 FRT/TO cells [53] with the Neon Transfection System (Invitrogen) following instructions from the supplier. Positively transfected cells (i.e. expressing the GFP reporter) were selected 48 hours after transfection with the cell sorter FACSAria (BD Biosciences) pooled together and maintained. Cells were then analyzed by IF using anti-AKAP450 antibody (A24), anti-CDK5Rap2 antibody (Millipore) and anti-PCNT (Abc-R), respectively in order to assess the efficiency of the mutagenesis process. Populations with a high frequency of mutant cells (i.e. more than 10% of cells negative for antibody staining) were used to isolate cells in a 96 well plate format to generate clones. Pure clones carrying the mutations were selected by immunofluorescence and further analyzed. In order to sequence the mutated genomic region a fragment of approximately 500 bp covering the targeted area was amplified by PCR with the following primers: *akap9-FW* 5′ AAGCAGTGAATGACAGTGCC, *akap9-RV* 5′ ATCCCTGTCAAAATCCGTGG; *cdk5rap2-FW* 5′ CTAGAAAAGCAAACACGAGG, *cdk5rap2-RV* 5′ TTGTCCAACTCTAGCCAAGG; *pcnt-FW* 5′ GCTCTGTTATCCCCACAGGGCACAG, *pcnt-RV* 5′ ACACCGTGACTTCCAGACACACAGG. The PCR products were cloned in pGEM-T vector and sequenced using SP6 and T7 primers.

### Western Blotting, Immunoprecipitation and MT-regrowth assays

For WB, proteins were separated by SDS/PAGE and gels were transferred to nitrocellulose filters and blocked for 1 h at 37°C in TBST (10mM Tris/HCl (pH7.4)/150mM NaCl/0.1% (v/v) Tween 20) containing 5% (w/v) non-fat dried milk. Filters were then incubated for 1-2h at 37°C with the primary antibody in the same buffer, washed and incubated for 45 min at 37°C with the secondary anti-rabbit or anti-mouse IgG antibodies conjugated with peroxidase (Amersham). For co-IPs, cells were lysed in NP40 buffer (10mM Tris/HCl (pH7.4)/150mM NaCl/10% (v/v) glycerol/ 1% (v/v) Nonidet P40/ 1mM PMSF and 1μg/ml of each pepstatin, leupeptin and aprotinin) for 20 min at 4°C and then for 3 min at 37°C. The extract was centrifuged at 20000g for 20 min and soluble fraction preadsorbed with an irrelevant antibody on Protein A-Sepharose or Protein G-agarose for 2h. Then extract was incubated with beads alone or immunoprecipitated with 0.5-1μl of the antibody of interest on Protein A-sepharose or Protein G-agarose for another 2 h. After washing beads pellets were analyzed by WB.

For repolymerization experiments, MTs were depolymerized with NZ (10 μM) for 3 h. Cells were rinsed five times with ice - cold medium and regrowth was induced by incubation in pre-warmed medium (37°C). All the MT regrowth experiments were carried out at room temperature.

### Immunofluorescence and image analysis

Cells were grown on glass coverslips and fixed with cold methanol (6 min at −20°C). Then, cells were incubated with primary antibodies for 1 h at RT, washed with PBS-Tween 0,1% and incubated with the appropriate fluorescent secondary antibody for 40 min. Nuclei were counterstained with DAPI (1 μg/ml) after secondary antibody labeling.

Confocal images were captured by a confocal Leica TCS SP5 using a HCX PL APO Lambda blue 63 × 1.4 OIL objective at 22°C and they correspond to maximal projections. Image processing was carried out using the Leica (LAS) and Adobe Photoshop softwares. For presentation, whole images were adjusted for intensity level, contrast and/or brightness.

For quantification of MT mass polymer, cell area and MT density (MT mass polymer/cell area), images were processed with Metamorph Offline software. Regions of interest were drawn around cells and after background subtraction, resulting fluorescence intensities and areas were estimated with the “region measurements” tool. For quantification of the fluorescence intensity of AKAP450, CDK5Rap2, PCNT and γ-tublin at the centrosome, a similar procedure was used but a ROI of 1.5 μm radius was drawn around the CAP350 signal.

Analysis of co-localization between the different markers and the golgi membranes was done by using MetaMorph software. All images from the same experiment were acquired using identical microscope settings and avoiding saturation of the brightest pixels. A threshold was set individually for each channel and applied to all the images. The software calculates the intensity of co-localization between the two channels where both labelings are positive over the threshold. Regions of interest were drawn in the merged image around cells excluding the centrosome and overlayed on the others channels. All the colocalization experiments were done in nocodazole-treated cells. For quantification of EB1 intensity at the Golgi membranes, microtubules were allowed to polimeryze for 3 min.

Quantification of EB1 comets in z-projected images was done in ImageJ as previously described [54]. Briefly, the background signal was subtracted using a rolling-ball radius of 50 and EB1 comets were detected with the “Find Maxima” tool. For quantification of EB1 comets around the centrosome, a ROI of 3 μm radius was drawn around the CAP350 signal at the centrosome. Determination of association between PCM proteins and Golgi fragments in control and centrinone-treated cells was performed by manually selecting those golgi fragments with visible associated PCM-protein foci and quantifying the total integration/fluorescence intensity of these spots, after background signal subtraction.

For quantifying MT nucleation from the GA, control and centrinone-treated cells were subjected to a 3-min MT regrowth assay, fixed and labeled for EB1 (green) and giantin (red). Double channel confocal images were analyzed by a customized software developed by Wimasis (Wimasis GmbH, Munich, Germany). For each image, the total number of growing microtubules, the number of MT-nucleating Golgi fragments and the number of MTs nucleated per Golgi fragment were determined. Finally, single cells were delineated and centrosomes of control cells were also identified and excluded from the analysis.

For analysis of MT nucleation from the GA in washout experiments, we select cells with either 6 or ≥8 visible centrioles and divide them into two categories: cells with or without MT-nucleation activity from the GA. Control and centrinone-treated cells were also analyzed for comparison.

### FACS analysis

For cell cycle analysis of control and centrinone-treated RPE-1 cells, cells were collected by centrifugation, washed with PBS and fixed in 70% ethanol at −20°C for at least 1h. After fixation, cells were washed twice with PBS and pellets were re-suspended in DNA staining solution (40 μg/mL propidium iodide (PI), 100 μg/mL RNase in PBS) for 30 minutes at 37°C in the dark. The DNA content was determined on a BD FACSCalibur flow cytometer (Becton Dickinson, USA). For cell size analysis of the same control and centrinone-treated RPE-1 cells, forward scatter (FSC-H) was used as a measure of the cell size.

### Statistical analysis

Quantitative data are expressed as mean ± SD. Alternatively, data values are displayed either in scatter or box-and-whisker plots. Significant differences among groups were evaluated by unpaired two-tailed Student’s t-test or one-way ANOVA followed by Tukey ot Dunnett’s multiple comparison tests, as appropriate (GraphPad Prism software) and indicated when relevant.

## ACKNOWLEDGMENTS

We thank K Oegema and A Siau (Ludwig Institute for Cancer Research, La Jolla, CA) for generously providing centrinone. We are grateful to L Pelletier (Lunenfeld-Tanenbaum Research Institute, Toronto, Canada) and Y Misumi (Fukuoka University, Japan) for antibodies. RPE-1 FRT/TO cells were a kind gift from J Pines (Gurdon Institute, Cambridge, UK). We also thank for the help provided by members of Microscopy Core Facility of CABIMER. This work was made possible by funding from the the Ministerio de Economía y Competitividad and Junta de Andalucía (grants BMC2012-36717, BFU2015-65747-P and CTS-2071 to RM Rios). P Gandolfo was supported by the Ministerio de Educación, Cultura y Deporte through a FPU predoctoral fellowship. FR Balestra is a recipient of a Marie Curie IEF postdoctoral fellowship.

## ABBREVIATIONS

Ab: Antibody
CTR: Centrosome
GA: Golgi apparatus
IF: Immunofluorescence
IP: Immunoprecipitation
MT: Microtubule
MTOC: Microtubule-organizing center
NZ: Nocodazole
PACT-domain: PCNT-AKAP450 centrosomal targeting
PCM: Pericentriolar material
KO: Knock-out
WB: Western blot

